# Distinct patterns of directed brain connectivity in focused attention, open monitoring and loving kindness meditation: An EEG Granger causality study with long-term meditators

**DOI:** 10.1101/2025.07.01.662572

**Authors:** Vasil Kolev, Kosio Beshkov, Peter Malinowski, Antonino Raffone, Juliana Yordanova

## Abstract

The present study applied spectral Granger causality analysis to electroencephalographic (EEG) recordings obtained during Focused Attention Meditation (FAM), Open Monitoring Meditation (OMM), and Loving Kindness Meditation (LKM) in highly experienced meditators. The aim of the investigation was to uncover distinct connectivity signatures associated with each meditation style by examining the strength, frequency band, and direction of inter-regional information transfers. These differences were expected to highlight the neural grounds of the cognitive and affective state of each meditative practice.

Multivariate Granger causality (GC) was computed from high-resolution EEG signals recorded from long-term meditators (n = 22) in four conditions: rest, FAM, OMM, and LKM. GC was analyzed in the frequency domain for key cortical regions (frontal and parietal) in the two hemispheres to compare frequency-specific directed connectivity between rest and each meditation type.

Main results demonstrated that each meditation state produced highly specific alterations in information transfer relative to rest. In FAM, there was significant reduction in posterior-to-anterior GC in the alpha and beta bands, and decreased multi-spectral inter-hemispheric frontal GC pointing to attenuated bottom-up sensory and associative inputs. In OMM, multi-spectral GC was significantly increased from the left hemisphere to the right posterior cortex implying expanded awareness in the right posterior regions through enhanced top-down modulation by the left-hemisphere. The distinctive features of LKM profile were the inter-hemispheric symmetry, the posterior-anterior bi-directionality, and the specific beta-band engagement, implying a co-activation of systems that support an emotionally balanced stance, equanimity and pro-social attitude.

These novel findings demonstrate that the direction and frequency specificity of information flows provide complementary insights into neural processes underlying distinct meditative states.

## 1. INTRODUCTION

Contemporary neuroscience increasingly recognizes that meditation does not modulate isolated brain regions in a uniform manner but rather orchestrates dynamic interactions within and between large-scale brain networks. Different meditative practices emphasize distinct cognitive and affective processes - such as sustained attention, meta-awareness, emotional regulation, and prosocial affect - and these processes are subserved by dissociable neural circuits.

In *Focused Attention Meditation* (FAM), practitioners actively sustain attention on a chosen object while inhibiting distractions. This practice has been associated with increased connectivity within the frontoparietal control network and dorsal attention network, supporting top-down attentional regulation and cognitive control (Tang et al., 2015; Fox et al., 2016). EEG studies have further shown elevated frontal midline theta and beta coherence, especially between frontal and parietal regions, indicating enhanced executive processing and sustained attentional effort (Cahn et al., 2010; Lomas et al., 2015; Yordanova et al., 2021).

In contrast, *Open Monitoring Meditation* (OMM) cultivates non-reactive awareness of moment-to-moment experience without focusing on a particular object. This style has been linked to more distributed and flexible brain dynamics, including decreased coupling between the striatum and posterior default mode network (DMN) regions such as the posterior cingulate cortex, suggesting diminished self-referential processing and increased cognitive flexibility (Lutz et al., 2008; Brewer & Garrison, 2014). EEG and MEG studies report increased gamma-band synchrony in right frontoparietal networks during OMM, reflecting heightened sensory awareness and adaptive monitoring (Lutz et al., 2004).

*Loving-Kindness Meditation* (LKM), focuses on cultivating prosocial emotions like compassion and empathy, has been shown to increase connectivity between medial prefrontal and inferior frontal cortices, as well as posterior DMN hubs including the precuneus (Garrison et al., 2014; Mascaro et al., 2013). Long-term LKM practice is also associated with enhanced integration across the DMN, salience, and social cognition networks, aligning with the emotional and interpersonal dimensions of the practice (Valk et al., 2017).

While these findings have shed light on the involvement of specific networks during different meditative states, less is known about the directionality and frequency-specific dynamics of information flow within and between these networks. In particular, the temporal causal relationships and oscillatory characteristics that underpin altered connectivity during meditation remain largely unexplored. Most neuroimaging and electrophysiological studies to date have relied on measures of undirected functional connectivity (e.g., correlation, coherence), which cannot determine how information flows across networks. Effective connectivity approaches - such as Granger causality (GC) - allow inferences about directionality of influence among neural sources, providing crucial insight into how top-down (e.g., prefrontal-to-posterior) or bottom-up (e.g., sensory-to-frontal) interactions are modulated by meditative state. This is particularly important given that distinct meditation styles may differentially rely on anticipatory control versus receptive awareness or empathic attunement. Moreover, neural communication is inherently spectrally structured: different frequency bands subserve different modes of brain function, such as theta for internal monitoring and memory processes, alpha for inhibition and internal attention, and beta for sensorimotor integration and cognitive control (Klimesch, 1999; Palva & Palva, 2007; Engel & Fries, 2010). Therefore, capturing the frequency specificity of directed interactions is essential for disentangling the neurophysiological mechanisms underlying meditative states. Despite the growing interest in meditation neuroscience, the joint exploration of directionality and frequency-specific connectivity across large-scale networks in different meditative practices remains limited. The present study aims to fill this gap by applying spectral Granger causality analysis to high-density EEG recordings obtained during FAM, OMM, and LKM in highly experienced meditators. We hypothesize that each meditation style will exhibit a distinct pattern of large-scale information flow characterized by specific directional pathways and dominant frequency bands. These differences are expected to reflect the cognitive and affective emphasis of each meditative practice, offering mechanistic insights into how distinct contemplative states reconfigure the brain’s functional architecture. By examining the *strength, frequency band*, and *direction* of inter-regional information transfers, we aim to uncover distinct causal connectivity signatures associated with each meditation style.

## 2. MATERIALS AND METHODS

### 2.1 Participants

The group of long-term meditators (LTM) consisted of twenty two healthy volunteers (mean age = 44.2 years, age range 26-70 years, 4 females). They were right-handed and did not report any history of movement disorders, neurological, psychiatric or somatic diseases.

These participants were monks, nuns and novice practitioners residing at Amaravati Buddhist Monastery, in Southern England, and at Santacittarama Monastery, in Central Italy. Practices at both monasteries are aligned with the Tai Forest Theravada Buddhist tradition which is now established, widely acknowledged and influential in the West. Participants practiced FAM (Śamatha), OMM (Vipassanā) and LKM (Metta) meditation forms in a balanced way in this tradition, often in integrated sessions, including silent meditation retreats (3 months per year). Meditation expertise is measured in hours taking into account both practice in the monastic tradition and practice before monastic life. In this tradition, the monks, nuns and novice practitioners typically practice two hours per day with the monastery community, with a regular intensification of practice during retreats (with several meditation sittings during the 3-month Winter retreat). As suggested by the abbots of the monasteries, we estimated participants with an average of 100 hours of practice per month during monastic life, with a balance of FAM, OMM and LKM facets of meditation. The lifetime duration of meditation practice of the participants was estimated as a mean value = 19358 hours (SE=3164), range 900-50600 hours.

The study had prior approval by the dedicated Research Ethics Committee at Sapienza University of Rome, Italy. All participants gave informed consent before participation according to the Declaration of Helsinki.

### 2.2 Experimental design

Participants had to perform a non-meditative rest condition and three meditation conditions: FAM, OMM and LKM. The switching between conditions was cued by voice. The instructions for the four conditions, which were written together with the abbot of Amaravati Monastery, the internationally recognized teacher Ajahn Amaro, were as follows: Rest: “Rest in a non-meditative relaxed state, without falling in sleep, while allowing any spontaneous thoughts and feelings to arise and unfold in the field of experience”. Focused attention (Śamatha) meditation: “Sustain the focus of attention on breath sensations, such as at the nostrils, noticing readily and with acceptance any arising distraction, such as on thoughts or stimuli, and in case of detected distraction, return readily and gently to focus attention on the breath sensations”. Open monitoring (awareness) meditation: “With an open receptive awareness, observe the contents of experience as they arise, change and fade from moment to moment, without restrictions or judgments – such contents including breath and body sensations, sensations arising from contact with external stimuli, feelings and thoughts”.

Loving kindness (Metta) meditation: “Generate and sustain metta, acceptance and friendliness towards yourself and the experience in the present moment, as well as towards any being, in any state or condition”.

### 2.3 Procedure

#### 2.3.1 EEG recordings

EEG was recorded with eyes closed in blocks of duration of approximately 2.5-3 minutes each, while the above mentioned conditions were repeated two times. In such a way, four blocks were repeated twice in the following order: REST, FAM, OMM, LKM. Thus, there were approximately 5-6 minutes (2 blocks x 2.5-3minutes) of total recording time for each condition. EEG was acquired by a mobile wireless system produced by Cognionics (https://www.cognionics.net/mobile-128) using an electrode cap with 64 active Ag/AgCl electrodes located in accordance with the extended international 10/20 system and referenced to linked mastoids. Electrode impedances were kept below 10 kOhm. EEG signals were collected at a sampling rate of 500Hz. For control of ocular artefacts, vertical and horizontal electro-oculogram (EOG) was also recorded. For experienced meditators, the experiment was conducted at the two monasteries in a quiet, dark room suitable for meditation and recording EEG. All participants were tested with the same experimental design.

#### 2.3.1 EEG pre-processing

EEG pre-processing was performed by means of Brain Vision Analyzer 2.2 (Brain Products GmbH, Germany). EEG analysis of Granger causality were conducted by means of software developed on Matlab R2013b (The Math Works Inc.).

EEG traces were visually inspected to reject epochs with noise or non-physiological artefacts. Bad channels were interpolated according to Hjorth (1975). All EEG traces were EOG corrected by means of independent component analysis (ICA, Makeig et al., 1997). EEG data were down-sampled to reduce off-line the recording frequency of 500 Hz to 250 Hz for data analysis.

To achieve a reference-free evaluation, all data analyses were performed after current source density (CSD) transform of the signals (Perrin et al., 1989; Nunez et al., 1997). The exact mathematical procedure is presented in detail in Perrin et al. (1989). The algorithm applies the spherical Laplace operator to the potential distribution on the surface of the head. The CSD transform replaces the potential at each electrode with the current source density of the electrical field calculated from all neighbour electrodes, thus eliminating the reference potential. When applied with dense electrode arrays (48–256 electrodes, 64 in the present study), this procedure provides excellent estimates of the bioelectric activity of the cortical surface (Nunez & Pilgreen, 1991). In the present analyses, to improve spatial resolution and reduce the volume conduction between electrodes, CSD was applied following with the parameters: order of splines = 10, maximal degree of Legendre polynomials = 4, lambda = 1E-7. Edge electrodes were excluded from all analyses, so that the number of channels was reduced. All analyses were carried out with CSD transformed data from 50 electrodes.

EEG recordings from all conditions were segmented in equal-sized non-overlapping epochs of 4.096-s duration. The average number of epochs for each condition/participant was 58 ±20). There were no statistically significant differences in the number of accepted trials between the two groups for each condition (REST, FAM, OMM, LKM).

### 2.4 Analysis of Granger causality (GC) in the frequency domain

To support the applied methodology, the formalism of Granger causality in the frequency domains is briefly presented in the following section. One of the main reasons that Granger causality has generated so much interest in neuroscience is the fact that it can be reformulated to estimate the connectivity between neural sources in specific frequency bands (Geweke, 1982, 1984). In order to calculate spectral causality it is not correct to apply band-pass filters to the data and then apply Granger causality. Instead, a Fourier transform is applied to the time lagged regression coefficients and then the frequency specific Granger causality is calculated according to the formula:

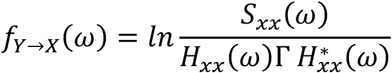

Here *S* represents the cross-power spectral density, *H* is the transformed (following Geweke, 1982) transfer function, with the star denoting the complex conjugate, and Γ is the residual covariance matrix. The full derivation of the frequency domain Granger causality is presented in detail in Chen et al. (2006) and Barnett & Seth (2014).

#### 2.4.1 GC computation pre-processing

Granger causality was analyzed using the publicly available MATLAB package (MVGC Matlab toolbox, Barnett & Seth, 2014). We paid particular attention to pre-processing steps given the sensitivity of GC to standard manipulations (Bressler & Seth, 2011; Seth, 2010). Analysis of Granger causality requires the analyzed signals to be stationary and not collinear (Seth, 2010). To avoid problems associated with these properties, the following additional pre-processing steps were undertaken.

First, as outlined before, a Laplacian filter (CSD) was applied to remove correlations in nearby electrodes, thereby also reducing the collinearity between signals. Second, following the requirements (Barnett & Seth, 2011, 2014), to remove mains-electricity line-noise (50 Hz) as well as its harmonic at 100 Hz which may lead to nonstationarity, notch filters (49–51 Hz and 99–101 Hz) were applied to the raw data. No other filtering was carried out. Also, since the length of the initially selected epochs in the data (4.096 s, 250 Hz) has been demonstrated to exhibit nonstationary features (e.g., Barrett et al., 2012), these epochs were additionally processed. They were divided into stationary non-overlapping segments of 1.024-s length. This length also was chosen to approach a balance between stationarity (shorter time series are more likely to be stationary) and model fit (longer time series support better parameter estimation for locally valid linear autoregressive models). This segmentation increased reliability by leading to a mean of 232±40 artefact-free EEG epochs per subject/per condition. After this step, the GC estimation still took an intractable amount of time. To ensure a reasonable model order for autoregressive modelling (Seth, 2010; Brovelli et al., 2004) and a reasonable amount of computation time, the selected artefact-free epochs were further subsampled by taking every fourth point from each time series. This effectively reduced the sampling rate by a factor of four (62.5 Hz). In this way, each epoch used for final analyses had a length of 1.024 s comprising 64 data points.

The third pre-processing step was to compute sensor-based regions of interest or electrode clusters (C) covering the left and right anterior and posterior cortical regions, for which specific connectivity patterns in meditation have been reported (Yordanova et al., 2020). We performed clustering as demonstrated in Fig. 1. The clustering scheme involved averaging the signals from the following electrodes: C1 (left anterior) = [F7, F5, F3, F1, FC5, FC3, FC1, C5, C3, C1]; C2 (right anterior) = [F8, F6, F4, F2, FC6, FC4, FC2, C6, C4, C2]; C3 (left posterior) = [CP5, CP3, CP1, P7, P5, P3, P1, PO7, PO3, O1]; C4 (right posterior) = [CP6, CP4, CP2, P8, P6, P4, P2, PO8, PO4, O2]. After the spatially averaged single-trial EEG of each defined cluster were computed, to avoid further confounds due to nonstationarity, a linear detrend was applied (Bernasconi & König, 1999; Hesse et al., 2003). Single-trial time-series corresponding to C1, C2, C3 and C4 were extracted for each subject and each condition and were used in subsequent analyses.

**Figure 1.**
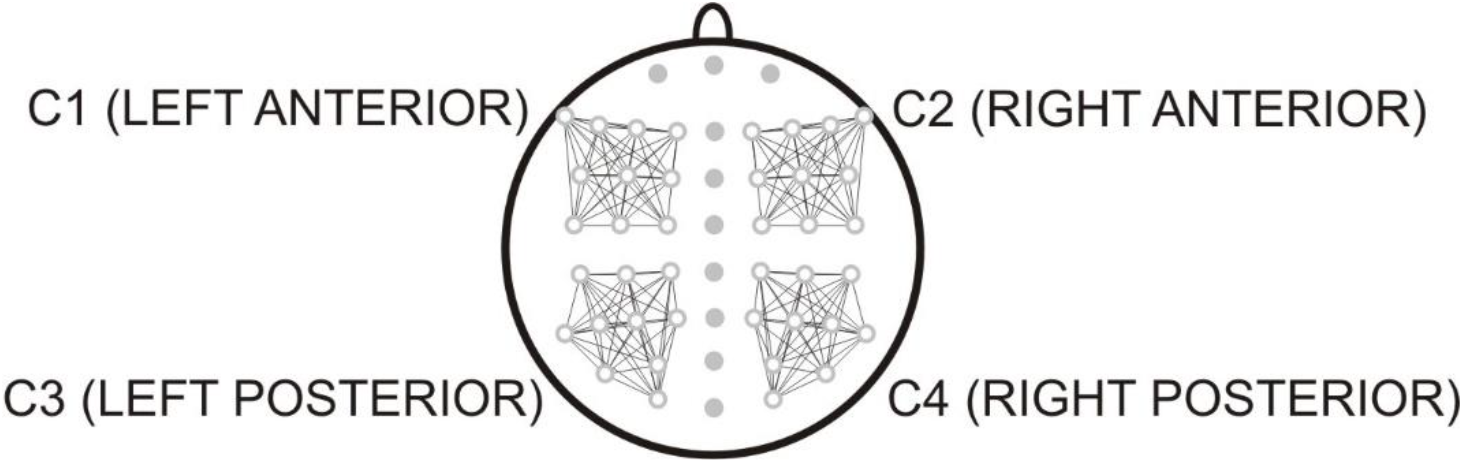
Schematic presentation of topographic clusters used for analysis: C1 (left anterior) = [F7, F5, F3, F1, FC5, FC3, FC1, C5, C3, C1]; C2 (right anterior) = [F8, F6, F4, F2, FC6, FC4, FC2, C6, C4, C2]; C3 (left posterior) = [CP5, CP3, CP1, P7, P5, P3, P1, PO7, PO3, O1]; C4 (right posterior) = [CP6, CP4, CP2, P8, P6, P4, P2, PO8, PO4, O2].

#### 2.4.2 Computation of GC

Computation of Granger causality was performed using the available MATLAB package MVGC Matlab toolbox (Barnett & Seth, 2014). This software was first applied to estimate the model order. For each single EEG epoch of the data, the recommended model order was computed as defined by the Akaike information criterion (AIC, Akaike, 1974; McQuarrie & Tsai, 1998). The 95th percentile of the values obtained was 10 (corresponding to 160 ms) and was used as model order throughout the GC analysis. The need of model order selection is to balance the number of parameters (as determined by the maximum order - lag) so as to achieve the best model fit to data while avoiding overfitting a finite data sequence. The next step was to maximise the likelihood function for the respective models. The MVGC toolbox facilities were used to obviate the need to estimate the reduced model parameters separately from the data. Specifically, to yield estimates asymptotically equivalent to the maximal likelihood estimate, ordinary least squares (OLS) were used (Hamilton, 1994). The estimated regression coefficients in the time domain were then transformed to an auto-covariance sequence through the Yule-Walker equations (Barnett & Sett, 2014), which was used to compute GC in the time domain. Finally, this sequence was used to calculate the Granger causalities in the frequency domain.

### 2.5 Statistical analysis

In order to compare the Granger causalities across frequencies between REST and FAM, OMM, and LKM, the following steps were followed: (1) After computing GC in the frequency domain, first, the Granger spectrum for each subject, each condition, each single trial, and each between-cluster pair was discretized into 1Hz bins. (2) Then, to compare the GC between REST and FAM, REST and OMM, and REST and LKM, all GC values within each 1Hz frequency bin for each trial/condition/pair in each group were concatenated and subjected to a non-parametric pair-wise Kruskal-Wallis test. Since this type of analysis involves multiple comparisons, the results were corrected with a false discovery rate (FDR) correction, as recommended for GC analysis in the frequency domain (Barnett & Seth, 2014).

## 3 RESULTS

Granger causality of experienced meditators was analyzed in the frequency domain with respect to delta, theta, alpha and beta frequency bands. Figure 2 depicts frequency-domain plots of GC for four clusters. A hertz-by-hertz FDR statistics applied to compare GC of each meditation state with GC during rest also is demonstrated in Fig. 2, and additionally summarized graphically in Fig. 3.

**Figure 2.**
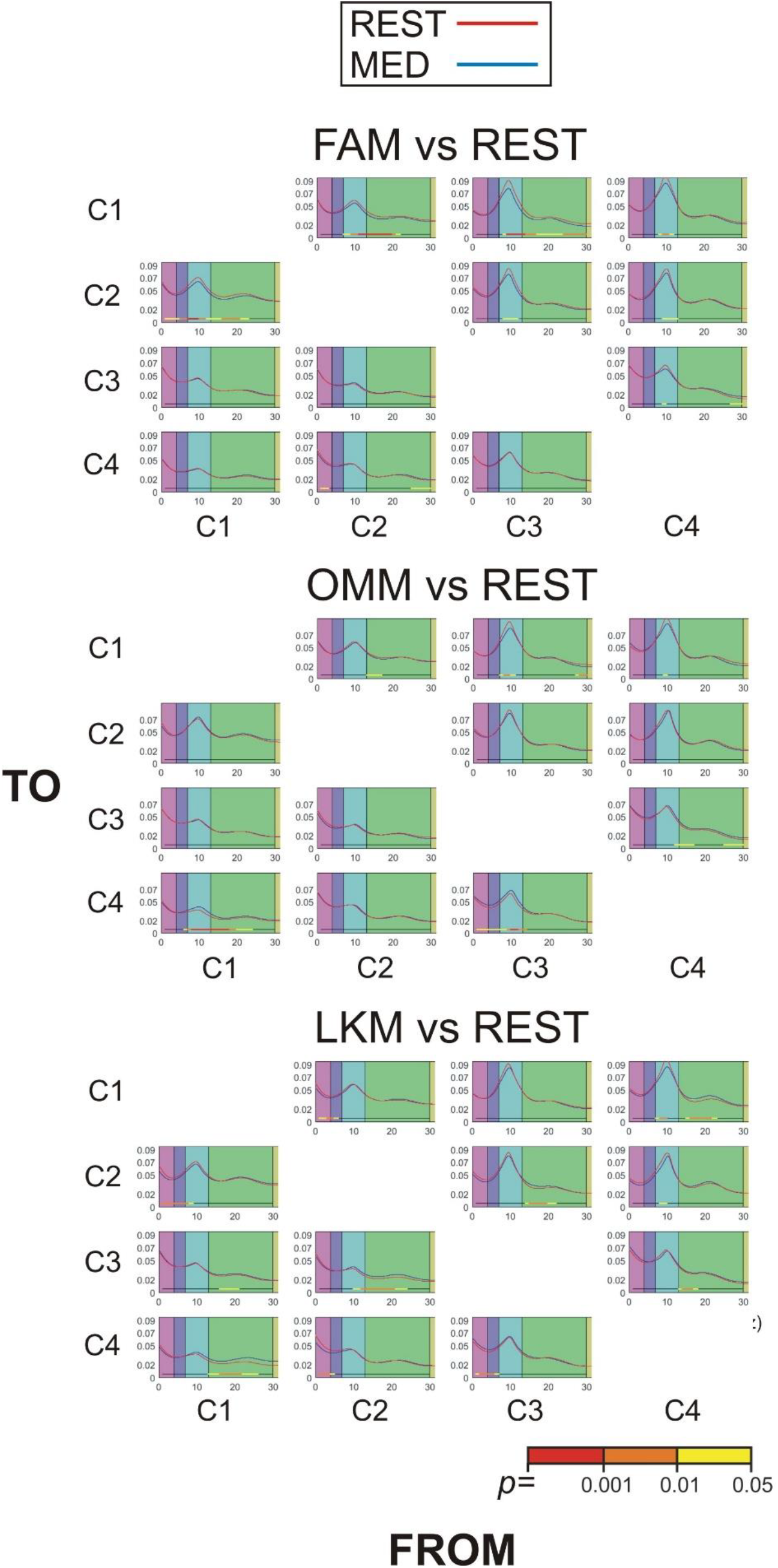
Frequency representation of Granger causality in long-term meditators: MEDITATION (MED) - blue line; REST - red line; FAM, Focused Attention Meditation; OMM, Open Monitoring Meditation; LKM, Loving Kindness Meditation.

**Figure 3.**
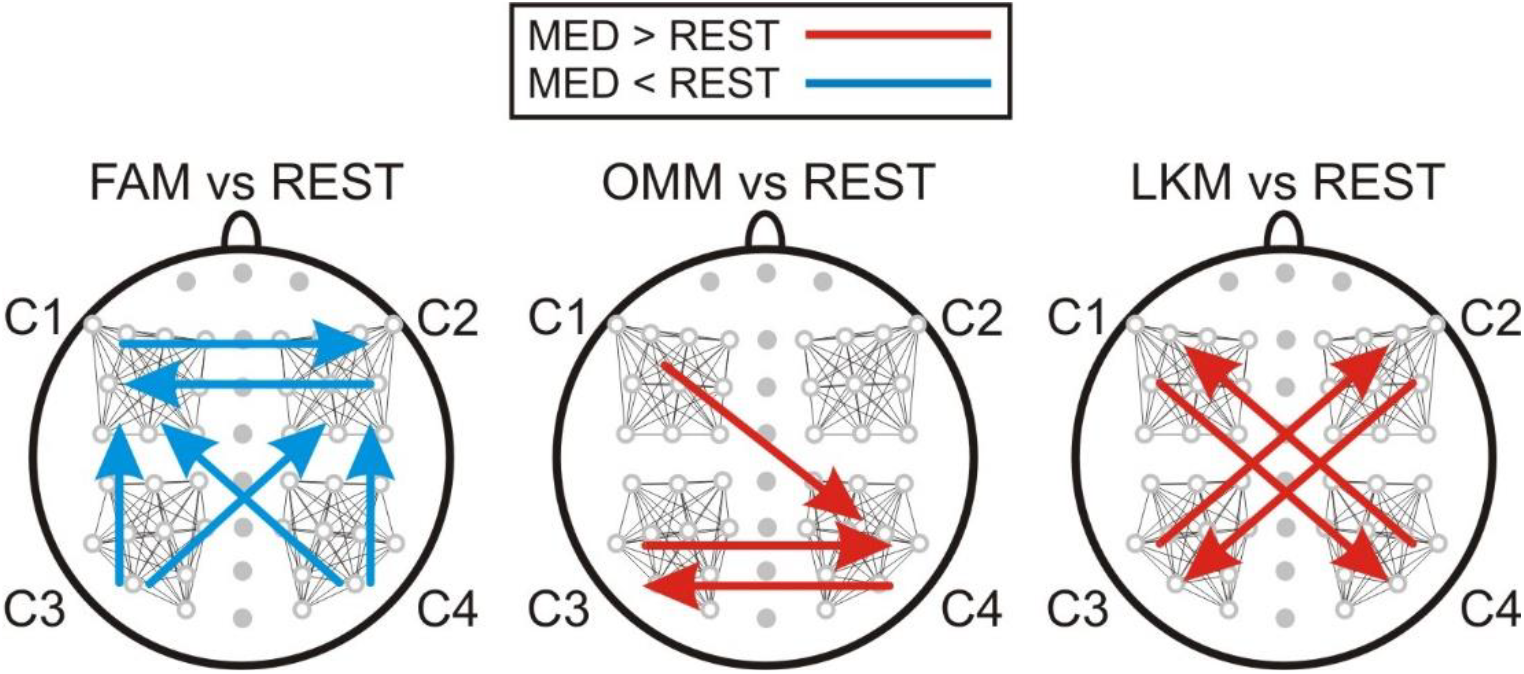
Schematic presentation of statistically significant differences between-conditions: MEDITATION (MED) > REST – red arrows; MED < REST – blue arrows; FDR corrected, p ≤ 0.05) in Granger causality in the frequency domain. Clusters: C1, C2, C3, C4. Focused Attention Meditation (FAM), Open monitoring Meditation (OMM), and Loving Kindness Meditation (LKM). Arrows indicate the direction of significantly different information flows.

Observation of frequency-domain representations of GC in Fig. 2 during rest and different types of meditation revealed the presence of a dominating broad peak in LTM with a maximum in the alpha band (8-10 Hz) encompassing also frequencies from the lower (theta) and higher (beta) bands. The broad alpha peak in LTM was most prominent for directed transfer connections from posterior (C3 and C4) to anterior (C1 and C2) clusters. A broad higher-frequency beta peak also was observed for most of the between-cluster connections.

*During FAM*, multi-spectral (delta/theta, alpha and beta) GC decreased bi-directionally for inter-hemispheric frontal connections. A suppression of mainly alpha directed flows was observed for both inter- and intra-hemispheric posterior-to-anterior connections (C3→C1, C2; C4→C1, C2). The posterior-to-anterior GC decrease was most expressed in the left hemisphere (C3→C1) including alpha and beta frequencies.

*During OMM*, the information flow from the left hemisphere to the right posterior region (C4) was significantly enhanced for multi-spectral GC (theta, alpha and beta). This was verified by the significant OMM-related GC increases for C1→C4 and C3→C4 (Figs. 2 and 3). The outflow from C4 was only enhanced to C3 in the beta band pointing to increased bi-directional communication between the posterior regions of the two hemispheres. In parallel, a slight decrease of the dominant posterior→anterior alpha peak was detected for intra-hemispheric connections.

*During LKM*, the main pattern of changes was characterized by a bi-directional cross-hemisphere increase of information flows between posterior and anterior regions supported by beta networks. These effects are demonstrated in Figs. 2 and 3 by the C1→C4, C4→C1, C2→C3, and C3→C2 GC increases in the beta band. The flows across posterior regions of the two hemispheres also increased in the slow (delta, theta) and fast (beta) frequency bands.

## 4 DISCUSSION

The present study employed EEG-based Granger causality analysis to investigate whether distinct meditation styles – FAM, OMM, and LKM - are associated with differences in the strength, EEG frequency, and direction of inter-regional information flow in highly experienced practitioners. It was hypothesized that the different activations and interactions of large-scale brain networks in specific meditative states (Brewer et al., 2011; Tang et al., 2015; Fox et al., 2014; Yordanova et al., 2021) would induce distinctive patterns of directed connectivity. The main results confirmed this hypothesis and demonstrated that each meditation state produced highly specific alterations in information transfer relative to rest, thus revealing novel neural signatures of meditation styles.

A deeper interpretation of state-specific modulations requires taking into account the fact that directed connectivity is altered in experienced meditators already during rest due to neuroplastic reorganization of their neural substrate. In a related study of GC (Kolev et al., in prep.), we compared the same sample of long-term meditators with novice meditators in resting state. We observed that extensive meditation practice produced a pronounced increase of information flow from posterior to frontal cortical regions and across frontal regions of the two hemispheres. The enhanced information transfer was multi-spectral involving theta, alpha and beta frequency bands, with most prominent expression of GC alpha peak. We interpreted the enhanced posterior-to-anterior multi-spectral transfer during rest as reflecting (1) increased bottom-up (sensory and interoceptive) signalling, and (2) neuroplastic modulation of fronto-parietal cognitive networks whereby posterior sensory and associative areas might inform rather than be directed by frontal executive regions. We proposed that meditation practice reorganizes the functioning of cognitive control by strengthening the major role of posterior regions in bottom-up attentional capture (Cabeza et al., 2008, 2012) and formation of “cross-modal and multi-modal integrative hubs” and “priority attentional maps” (Gottlieb et al., 2009; Humphreys et al., 2017). We suggested that an optimized formation of stable priority maps in long-term meditators facilitates an integrated multi-modal sensory input and minimizes frontal guidance for an effortless maintenance of mental states (Kolev et al., in prep.).

According to the present results, in the FAM condition, experienced meditators exhibited a significant reduction in posterior-to-anterior GC in the alpha and beta bands compared to resting state, alongside decreased multi-spectral inter-hemispheric frontal GC. In view of rest-related modulations (Kolev et al., in prep.) the reduction in posterior-to-anterior information transfer may reflect a downregulation of bottom-up sensory and associative inputs, aligning with the “gating by inhibition” model of alpha oscillations (Palva & Palva, 2007). By attenuating sensory-driven signalling from posterior regions, the brain may enhance internal stability and reduce perceptual interference. This is consistent with the cognitive demands of FAM, which emphasizes sustained, narrow attentional focus on a chosen object (e.g., the breath) while minimizing the influence of distractors (Cahn and Polich, 2006; Lutz et al., 2008; Malinowski et al., 2013). Similarly, the reduction in bilateral frontal GC suggests diminished demands for inter-hemispheric executive coordination.

Theta- and alpha-band frontal connectivity is typically associated with cognitive control and control allocation (Cavanagh & Frank, 2014; Cohen, 2011; Clayton et al., 2015). Hence, the selective suppression of information transfer across frontal regions is possibly due to the decreased need for cognitive flexibility, monitoring or task switching during the maintenance of focused attention. These reductions in information flows underscore the selective inhibition characteristics of advanced FAM practice, marking a shift from more distributed cortical communication toward local, functionally specialized loops.

During OMM, experienced meditators showed significantly enhanced directed connectivity from the left anterior to the right posterior cortex, as well as from the left to right posterior regions, across theta, alpha, and beta frequency bands. This pattern demonstrates an overall increase of directed influence of the left hemisphere over right posterior regions. This left-to-right posterior inflow may reflect several interwoven functional mechanisms that are specifically cultivated by OMM. ***First***, attentional fronto-parietal networks can be considered (Uddin et al., 2019). The bi-lateral dorsal attentional networks are associated with goal-directed sustained focus and top-down regulation of spatial and feature-based attention (Corbetta & Shulman, 2002; Szczepanski et al., 2013; Gross et al., 2004; Popov et al., 2017). In contrast, the right-lateralized ventral attention network including the right temporo-parietal junction and right inferior frontal gyrus is known to support stimulus-driven reorienting, especially to salient, novel, or distracting stimuli. Its function is crucial for detecting mind-wandering, internal distractions, or unexpected sensory inputs (Corbetta & Shulman, 2002; Vossel et al., 2014; Uddin et al., 2019). The major operating rhythms of attentional networks encompass both slow and fast-frequency EEG ranges, in particular theta, alpha and beta frequency bands (Daitch et al., 2013; Sadaghiani & Kleinschmidt, 2016; Yordanova et al., 2017). The asymmetric left-to-right posterior information transfer observed during OMM may reflect the volitional control of open attentional focus allowing for early detection and expanded monitoring of all external and internal mental events, as well as flexible dynamic shifts in attention as emphasized in OMM (Cahn & Polich, 2006; Lutz et al., 2008). ***Second,***the increased inflow to right posterior cortical regions also may reflect enhanced functional engagement of the right parietal cortex which is critical for awareness. Clinical evidence from patients with spatial neglect following right parietal damage verifies this region’s role in maintaining perceptual awareness of contralateral space (Vallar & Calzolari, 2018; Corbetta & Shulman, 2011; Yordanova et al., 2017). Also the right parietal cortex, particularly the inferior parietal lobule and temporo-parietal junction, is central to constructing the embodied sense of self in space and time, as demonstrated by neuroimaging studies showing the links to core aspects of bodily self-consciousness, including self-location, ownership, and agency (Blanke & Arzy, 2005; Blanke et al., 2015; Uddin et al., 2016; Convento et al., 2018; Ionta et al., 2014). Notably, the right temporo-parietal junction has also been implicated in interoceptive awareness (Critchley et al., 2004 Raimo et al., 2020. These findings suggest that the enhanced information transfer toward right parietal regions during OMM may underlie expanded awareness, consistent with mental experiences during OMM. ***Third,***the left-right posterior profile in OMM may additionally result from a stronger involvement of the left hemisphere in emotion processing and social cognition (Davidson, 1992), aligning with the non-reactive mode of acceptance and equanimity adopted in OMM. The strengthened posterior-to-posterior cross-hemispheric connectivity in theta and alpha bands may support integrative awareness by aligning bilateral perceptual fields and fostering coherent sensory processing without fixation (Chica et al., 2013). Together, these findings suggest that OMM in highly trained individuals promotes a neural architecture that enables expanded awareness and open receptivity through especially targeting right posterior cortices and enhancing its top-down modulation by the left-hemisphere. Such a configuration aligns with the cognitive requirements of OMM involving sustained monitoring of internal and external experiences without effortful selection or suppression, and expanding the scope of awareness beyond the focus typical of goal-directed tasks. Also, it may be suggested that previously identified predominant engagement of the left hemisphere across meditation styles and in long-term meditators (Fox et al., 2014; Marzetti et al., 2014; Guidotti et al., 2023; Yordanova et al., 2020, 2021; De Felippi et al., 2022; Raffone et al., 2019) may be essentially linked to faculties of mindfulness that are trained in OMM.

The present findings reveal a striking distinctive pattern of inter-hemispheric beta connectivity during LKM, specifically involving left frontal-right posterior and right frontal-left posterior regions. The distinctive features of this profile are the inter-hemispheric symmetry, the posterior-anterior bi-directionality, and the specific beta-band engagement.

The symmetric posterior-anterior transfer generally suggests enhanced large-scale integration across networks during compassion-based meditation. LKM involves the intentional cultivation of affective states such as compassion, kindness, and warmth toward self and others. This process may engage both prefrontal regions implicated in emotion regulation and cognitive control (Ochsner & Gross, 2005), and posterior regions (e.g., temporo-parietal junction, posterior cingulate cortex) implicated in self-referential processing, perspective-taking, and social cognition (Decety & Lamm, 2007; Lutz et al., 2008). Hence, the observed increase in GC between these regions likely reflects top-down modulation of affective and self-related representations, supporting the sustained generation of compassionate mental states. The detected symmetry of cross-hemispheric GC pattern (left frontal → right posterior and right frontal → left posterior) may be referred to the known lateralization of affective processing (Davidson, 2004). It may indicate a balanced recruitment of emotional approach and avoidance systems. The left frontal cortex is frequently associated with approach motivation and positive affect, whereas the right frontal cortex is linked to vigilance and withdrawal (Davidson, 1992; 2004). During LKM, the co-activation and coordination of these systems may underlie an emotionally balanced stance supporting equanimity and open-heartedness. Such symmetry may therefore be specifically critical for the generation of empathic resonance without emotional reactivity (Singer & Klimecki, 2014; Klimecki et al., 2014). This interpretation aligns with neuroimaging studies showing that LKM engages bilateral networks, including medial prefrontal cortex, anterior cingulate, insula, and temporo-parietal regions (Lutz et al., 2009; Mascaro et al., 2013). Another conspicuous observation in LKM was the specific engagement of directed beta connectivity, in contrast to the directed multi-spectral connectivity in FAM and OMM. Beta-band connectivity has been implicated in the maintenance of task goals and internal set, and coordination of top-down attention (Buschman & Miller, 2007; Cavanagh et al., 2009; Fries, 2015; Gaillard et al., 2009; Engel & Fries, 2010; Spitzer & Haegens, 2017). In LKM, practitioners aim to sustain an internal positive affective intention toward self and others. The observed frontal-posterior beta GC may thus reflect top-down entrainment of posterior associative regions to support a stable, compassionate mindset over time. This top-down control is needed to support a stabilized cognitive-affective mode, in which emotional intentions are maintained and integrated with sensory and social information. This view is supported by prior EEG studies of meditation showing enhanced beta coherence during concentrative and affective practices (Cahn et al., 2010), and by intracranial and MEG findings that link beta rhythms to intentional state maintenance and cross-modal integration (Bastos et al., 2015). These original observations of inter-hemispheric symmetric fronto-posterior pathways supported by beta networks in LKM might reveal novel correlates of perspective-taking and empathy-related processes in the brain. It is intriguing to suggest that the refined top-down control of mental states (emotional, contemplative) mainly relies on fast-frequency oscillatory networks, in contrast to the top-down control of object-based or event-based goals engaging theta/alpha-frequency networks.

One important issue that needs to be emphasized is that when the frequency characteristics of functional (undirected) connectivity were analyzed in the same sample of experienced meditators in the same meditative conditions – FAM, OMM, and LKM (Yordanova et al., 2020), connectivity patterns were revealed that were common for the three meditation styles. Specifically, common hubs were detected in the left posterior cortex supported by theta networks and in the right posterior cortex supported by alpha networks. Only a lateralization of undirected connectivity in the beta band distinguished the three meditation conditions.

These previous observations of undirected connectivity are in stark contrast with current findings on directed connectivity indicating conspicuously distinctive profiles in the three meditation states as compared to rest. They confirm that the direction and frequency specificity of information flows carry relevant additional information about underlying neural processes (Pullon et al., 2017; Hillebrand et al., 2016) that may provide complementary insights to understanding brain states.

## Acknowledgments

We would like to express our gratitude and appreciation to the monks, nuns and novices of Amaravati and Santacittarama Buddhist Monasteries for their outstanding dedication and participation in our study. This work has been supported by a grant from BIAL Foundation (Portugal) for the project “Advancements on the aware mind-brain: New insights about the neural correlates of meditation states and traits” (Grant 272/70), and by the National Research Fund by the Ministry of Education and Science, Sofia, Bulgaria (Project KP-06-N33/11/2019).

## Data availability

The datasets used and analyzed during the current study are available from the corresponding author on reasonable request.

## References

1. Akaike, H. (1974). A new look at the statistical model identification. IEEE Trans Autom Control, 19, 716–723. 10.1109/TAC.1974.1100705

2. Barnett, L., & Seth, A.K. (2014). The MVGC multivariate Granger causality toolbox: a new approach to Granger-causal inference. J Neurosci Methods, 223, 50–68. 10.1016/j.jneumeth.2013.10.018

3. Barrett, A.B., Murphy, M., Bruno, M.A., Noirhomme, Q., Boly, M., Laureys, S., & Seth, A.K. (2012). Granger causality analysis of steady-state electroencephalographic signals during propofol-induced anaesthesia. PLoS One, 7, e29072. 10.1371/journal.pone.0029072

4. Bastos, A.M., Vezoli, J., Bosman, C.A., Schoffelen, J.M., Oostenveld, R., Dowdall, J.R., De Weerd, P., Kennedy, H., & Fries, P. (2015). Visual areas exert feedforward and feedback influences through distinct frequency channels. Neuron, 85, 390–401. 10.1016/j.neuron.2014.12.018

5. Bernasconi, C., & Konig, P. (1999). On the directionality of cortical interactions studied by structural analysis of electrophysiological recordings. Biol Cybern, 81, 199–210. 10.1007/s004220050556

6. Blanke, O., & Arzy, S. (2005). The out-of-body experience: disturbed self-processing at the temporo-parietal junction. Neuroscientist, 11. 16–24. 10.1177/1073858404270885

7. Blanke, O., Slater, M., & Serino, A. (2015). Behavioral, neural, and computational principles of bodily self-consciousness. Neuron, 88, 145–66. 10.1016/j.neuron.2015.09.029

8. Brewer, J.A., & Garrison, K.A. (2014). The posterior cingulate cortex as a plausible mechanistic target of meditation: findings from neuroimaging. Ann. NY Acad. Sci., 1307, 19–27. 10.1111/nyas.12246

9. Brewer, J.A., Worhunsky, P.D., Gray, J.R., Tang, Y.Y., Weber, J., & Kober, H. (2011). Meditation experience is associated with differences in default mode network activity and connectivity. Proc Natl Acad Sci USA, 108, 20254–20259. 10.1073/pnas.1112029108

10. Brovelli, A., Ding, M., Ledberg, A., Chen, Y., Nakamura, R., & Bressler, S. (2004). Beta oscillations in a largescale sensorimotor cortical network: Directional influences revealed by Granger causality. Proc Natl Acad Sci USA, 101, 9849–9854. 10.1073/pnas.0308538101

11. Buschman, T.J., & Miller, E.K. (2007). Top-down versus bottom-up control of attention in the prefrontal and posterior parietal cortices. Science, 315, 1860–1864. 10.1126/science.1138071

12. Cabeza, R., Ciaramelli, E., & Moscovitch, M. (2012). Cognitive contributions of the ventral parietal cortex: an integrative theoretical account. Trends Cogn Sci, 16, 338–352. 10.1016/j.tics.2012.04.008

13. Cabeza, R., Ciaramelli, E., Olson, I.R., & Moscovitch, M. (2008). The parietal cortex and episodic memory: an attentional account. Nat Rev Neurosci, 9, 613–625. 10.1038/nrn2459

14. Cahn, B.R., Delorme, A., & Polich, J. (2010). Occipital gamma activation during Vipassana meditation. Cognitive Processing, 11, 39–56. 10.1007/s10339-009-0352-1

15. Cahn, B.R., & Polich, J. (2006). Meditation states and traits: EEG, ERP, and neuroimaging studies. Psychol Bull, 132, 180–211. 10.1037/0033-2909.132.2.180

16. Cavanagh, J.F., Cohen, M.X., & Allen, J.J. (2009). Prelude to and resolution of an error: EEG phase synchrony reveals cognitive control dynamics during action monitoring. J Neurosci, 29, 98–105. 10.1523/JNEUROSCI.4137-08.2009

17. Cavanagh, J.F., & Frank, M.J. (2014). Frontal theta as a mechanism for cognitive control. Trends Cogn Sci, 18, 414–421. 10.1016/j.tics.2014.04.012

18. Chen, Y., Bressler, S.L., & Ding, M. (2006) Frequency decomposition of conditional Granger causality and application to multivariate neural field potential data. J Neurosci Methods, 150, 228–237. 10.1016/j.jneumeth.2005.06.011

19. Chica, A.B., Paz-Alonso, P.M., Valero-Cabré, A., & Bartolomeo, P. (2013). Neural bases of the interactions between spatial attention and conscious perception. Cereb Cortex, 23, 1269–1279. 10.1093/cercor/bhs087

20. Clayton, M.S., Yeung, N., & Kadosh, R.C. (2015). The roles of cortical oscillations in sustained attention. Trends Cogn Sci, 19, 188–195. 10.1016/j.tics.2015.02.004

21. Cohen, M. X. (2011). Error-related medial frontal theta activity predicts cingulate-related structural connectivity. NeuroImage, 55, 1373–1383. 10.1016/j.neuroimage.2010.12.072

22. Convento, S., Romano, D., Maravita, A., & Bolognini, N. (2018). Roles of the right temporo-parietal and premotor cortices in self-location and body ownership. Eur J Neurosci, 47, 1289–1302. 10.1111/ejn.13937

23. Corbetta, M., & Shulman, G.L. (2002). Control of goal-directed and stimulus-driven attention in the brain. Nat Rev Neurosci, 3, 201–215. 10.1038/nrn755

24. Corbetta, M., & Shulman, G.L. (2011). Spatial neglect and attention networks. Annu Rev Neurosci, 34, 569–599. 10.1146/annurev-neuro-061010-113731

25. Critchley, H.D., Wiens, S., Rotshtein, P., Ohman, A., & Dolan, R.J. (2004). Neural systems supporting interoceptive awareness. Nat Neurosci, 7, 189–195. 10.1038/nn1176

26. Daitch A.L., Sharma M., Roland J.L., Astafiev S.V., Bundy D.T., Gaona C.M., Snyder A.Z., Shulman G.L., Leuthardt E.C., & Corbetta M. (2013). Frequency-specifc mechanism links human brain networks for spatial attention. Proc. Natl. Acad. Sci. USA, 110, 19585–19590. 10.1073/pnas.1307947110

27. Davidson, R.J. (1992). Anterior cerebral asymmetry and the nature of emotion. Brain Cogn, 20, 125–151. 10.1016/0278-2626(92)90065-t

28. Davidson, R.J. (2004). What does the prefrontal cortex “do” in affect: perspectives on frontal EEG asymmetry research. Biol Psychol, 67, 219–233. 10.1016/j.biopsycho.2004.03.008

29. De Filippi, E., Escrichs, A., Càmara, E., Garrido, C., Marins, T., Sánchez-Fibla, M., Gilson, M., & Deco, G. (2022). Meditation-induced effects on whole-brain structural and effective connectivity. Brain Struct Funct, 227, 2087–2102. 10.1007/s00429-022-02496-9

30. Decety, J., & Lamm, C. (2007). The role of the right temporoparietal junction in social interaction: how low-level computational processes contribute to meta-cognition. Neuroscientist, 13, 580–593. 10.1177/1073858407304654

31. Engel, A.K., & Fries, P. (2010). Beta-band oscillations--signalling the status quo? Curr Opin Neurobiol, 20, 156–165. 10.1016/j.conb.2010.02.015

32. Fox, K.C.R., Dixon, M.L., Nijeboer, S., Girn, M., Floman, J.L., Lifshitz, M., Ellamil, M., Sedlmeier, P., & Christoff, K. (2016). Functional neuroanatomy of meditation: A review and meta-analysis of 78 functional neuroimaging investigations. Neuroscience & Biobehavioral Reviews, 65, 208–228. 10.1016/j.neubiorev.2016.03.021

33. Fox, K.C.R., Nijeboer, S., Dixon, M.L., Floman, J.L., Ellamil, M., Rumak, S.P., Sedlmeier, P., & Christoff, K. (2014). Is meditation associated with altered brain structure? A systematic review and meta-analysis of morphometric neuroimaging in meditation practitioners. Neurosci. Biobehav. Rev., 43, 48–73. 10.1016/j.neubiorev.2014.03.016

34. Fries, P. (2015). Rhythms for cognition: Communication through coherence. Neuron, 88, 220–235. 10.1016/j.neuron.2015.09.034

35. Gaillard, R., Dehaene, S., Adam, C., Clémenceau, S., Hasboun, D., Baulac, M., Cohen, L., & Naccache, L. (2009). Converging intracranial markers of conscious access. PLoS Biol, 7, e61. 10.1371/journal.pbio.1000061

36. Garrison, K.A., Zeffiro, T.A., Scheinost, D., Constable, R.T., & Brewer, J.A. (2014). Meditation leads to reduced default mode network activity beyond an active task. Cognitive, Affective, & Behavioral Neuroscience, 15, 712–720. 10.3758/s13415-015-0358-3

37. Geweke, J.F. (1982). Measurement of linear dependence and feedback between multiple time series. J. Am. Stat. Assoc., 77, 304–313.

38. Geweke J.F. (1984). Measures of conditional linear dependence and feedback between time series. J. Am. Stat. Assoc., 79, 907–915.

39. Gottlieb, J., Balan, P., Oristaglio, J., & Suzuki, M. (2009). Parietal control of attentional guidance: the significance of sensory, motivational and motor factors. Neurobiol Learn Mem, 91, 121–128. 10.1016/j.nlm.2008.09.013

40. Gross, J., Schmitz, F., Schnitzler, I., Kessler, K., Shapiro, K., Hommel, B., & Schnitzler A. (2004). Modulation of long-range neural synchrony reflects temporal limitations of visual attention in humans. Proc. Natl. Acad. Sci. USA, 101, 13050–13055. 10.1073/pnas.0404944101

41. Guidotti, R., D’Andrea, A., Basti, A., Raffone, A., Pizzella, V., & Marzetti, L. (2023). Long-term and meditation-specific modulations of brain connectivity revealed through multivariate pattern analysis. Brain Topogr, 36, 409–418. 10.1007/s10548-023-00950-3

42. Hamilton, J.D. (1994). Time Series Analysis. Princeton: Princeton University Press.

43. Hesse, W., Moeller, E., Arnold, M., & Schack, B. (2003). The use of time-variant EEG Granger causality for inspecting directed interdependencies of neural assemblies. J Neurosci Meth, 124, 27–44. 10.1016/s0165-0270(02)00366-7

44. Hillebrand, A., Tewarie, P., van Dellen, E., Yu, M., Carbo, E.W., Douw, L., Gouw, A.A., van Straaten, E.C., & Stam, C.J. (2016). Direction of information flow in large-scale resting-state networks is frequency-dependent. Proc Natl Acad Sci USA, 113, 3867–3872. 10.1073/pnas.1515657113

45. Hjorth, B. (1975). An on-line transformation of EEG scalp potentials into orthogonal source derivations. Electroencephalogr. Clin. Neurophysiol., 39, 526–530. 10.1016/0013-4694(75)90056-5

46. Humphreys, G.F., & Lambon Ralph, M.A. (2017). Mapping domain-selective and counterpointed domain-general higher cognitive functions in the lateral parietal cortex: Evidence from fMRI comparisons of difficulty-varying semantic versus visuo-spatial tasks, and functional connectivity analyses. Cereb. Cortex, 27, 4199–4212. 10.1093/cercor/bhx107

47. Ionta, S., Martuzzi, R., Salomon, R., & Blanke, O. (2014). The brain network reflecting bodily self-consciousness: a functional connectivity study. Soc Cogn Affect Neurosci, 9, 1904–1913. 10.1093/scan/nst185

48. Klimecki, O.M., Leiberg, S., Ricard, M., & Singer, T. (2014). Differential pattern of functional brain plasticity after compassion and empathy training. Soc Cogn Affect Neurosci, 9, 873–879. 10.1093/scan/nst060

49. Klimesch, W. (1999). EEG alpha and theta oscillations reflect cognitive and memory performance: a review and analysis. Brain Res Rev, 29, 169–195. 10.1016/s0165-0173(98)00056-3

50. Kolev, V., Beshkov, K., Malinowski, P., Raffone, A., & Yordanova, J. Neuroplasticity of directed connectivity in long-term meditation: Evidence from EEG Granger causality, in preparation.

51. Lomas, T., Ivtzan, I., & Fu, C.H.Y. (2015). A systematic review of the neurophysiology of mindfulness on EEG oscillations. Neuroscience & Biobehavioral Reviews, 57, 401–410. 10.1016/j.neubiorev.2015.09.018

52. Lutz, A., Greischar, L.L., Perlman, D.M., & Davidson, R.J. (2009). BOLD signal in insula is differentially related to cardiac function during compassion meditation in experts vs. novices. NeuroImage, 47, 1038–1046. 10.1016/j.neuroimage.2009.04.081

53. Lutz, A., Greischar, L.L., Rawlings, N.B., Ricard, M., & Davidson, R.J. (2004). Long-term meditators self-induce high-amplitude gamma synchrony during mental practice. Proceedings of the National Academy of Sciences, 101, 16369–16373. 10.1073/pnas.0407401101

54. Lutz, A., Slagter, H.A., Dunne, J.D., & Davidson, R.J. (2008). Attention regulation and monitoring in meditation. Trends in Cognitive Sciences, 12, 163–169. 10.1016/j.tics.2008.01.005

55. Makeig, S., Bell, A.J., Jung, T.-P., Ghahremani, D., & Sejnowski, T.J. (1997). Blind separation of auditory event-related brain responses into independent components. Proc Natl Acad Sci USA, 94, 10979–10984, 10.1073/pnas.94.20.10979

56. Malinowski, P. (2013). Neural mechanisms of attentional control in mindfulness meditation. Front. Neurosci., 7, 8. 10.3389/fnins.2013.00008

57. Marzetti, L., Di Lanzo, C., Zappasodi, F., Chella, F., Raffone, A., & Pizzella, V. (2014). Magnetoencephalographic alpha band connectivity reveals differential default mode network interactions during focused attention and open monitoring meditation. Front. Hum. Neurosci., 8, 832. 10.3389/fnhum.2014.00832

58. Mascaro, J.S., Rilling, J.K., Negi, L.T., & Raison, C.L. (2013). Pre-existing brain function predicts subsequent practice of mindfulness and compassion meditation. NeuroImage, 69, 35–42. 10.1016/j.neuroimage.2012.12.021

59. McQuarrie, A.D.R., & Tsai, C.L. (1998). Regression and Time Series Model Selection. Singapore: World Scientific Publishing.

60. Nunez, P.L., & Pilgreen, K.L. (1991). The spline-Laplacian in clinical neurophysiology: a method to improve EEG spatial resolution. J. Clin. Neurophysiol., 8, 397–413.

61. Nunez, P.L., Srinivasan, R., Westdorp, A.F., Wijesinghe, R.S., Tucker, D.M., Silberstein, R.B., & Cadusch, P.J. (1997). EEG coherency. I. Statistics, reference electrode, volume conduction, Laplacians, cortical imaging, and interpretation at multiple scales. Electroencephalogr. Clin. Neurophysiol., 103, 499–515. 10.1016/s0013-4694(97)00066-7

62. Ochsner, K.N., & Gross, J.J. (2005). The cognitive control of emotion. Trends Cogn Sci, 9, 242–249. 10.1016/j.tics.2005.03.010

63. Palva, S., & Palva, J.M. (2007). New vistas for alpha-frequency band oscillations. Trends Neurosci, 30, 150–158. 10.1016/j.tins.2007.02.001

64. Perrin, F., Pernier, J., Bertrand, O., & Echallier, J.F. (1989). Spherical splines for scalp potential and current density mapping. Electroencephalogr. Clin. Neurophysiol., 72, 184–187. 10.1016/0013-4694(89)90180-6

65. Popov, T., Kastner, S., & Jensen, O. (2017). FEF-controlled alpha delay activity precedes stimulus-induced gamma-band activity in visual cortex. J Neurosci, 37, 4117–4127. 10.1523/JNEUROSCI.3015-16.2017

66. Pullon, R.M., Yan, L., Sleigh, J.W., & Warnaby, C.E. (2020). Granger causality of the electroencephalogram reveals abrupt global loss of cortical information flow during propofol-induced loss of responsiveness. Anesthesiology, 133, 774–786. 10.1097/ALN.0000000000003398

67. Raffone, A., Marzetti, L., Del Gratta, C., Perrucci, M.G., Romani, G.L., & Pizzella, V. (2019). Toward a brain theory of meditation. Prog Brain Res, 244, 207–232. 10.1016/bs.pbr.2018.10.028

68. Raimo, S., Boccia, M., Di Vita, A., Iona, T., Cropano, M., Ammendolia, A., Colao, R., Iocco, M., Angelillo, V., Guariglia, C., Grossi, D., & Palermo, L. (2020). Interoceptive awareness in focal brain-damaged patients. Neurol Sci, 41, 1627–1631. 10.1007/s10072-019-04172-z

69. Sadaghiani, S., & Kleinschmidt, A. (2016). Brain networks and α-oscillations: Structural and functional foundations of cognitive control. Trends Cogn Sci, 20, 805–817. 10.1016/j.tics.2016.09.004

70. Seth, A.K. (2010). A MATLAB toolbox for Granger causal connectivity analysis. Neurosci Meth, 186, 262–273. 10.1016/j.jneumeth.2009.11.020

71. Singer, T., & Klimecki, O.M. (2014). Empathy and compassion. Curr Biol, 24, R875–R878. 10.1016/j.cub.2014.06.054

72. Spitzer, B., & Haegens, S. (2017). Beyond the status quo: A role for beta oscillations in endogenous content (re)activation. eNeuro, 4, ENEURO.0170-17.2017. 10.1523/ENEURO.0170-17.2017

73. Szczepanski, S.M., Pinsk, M.A., Douglas, M.M., Kastner, S., & Saalmann, Y.B. (2013). Functional and structural architecture of the human dorsal frontoparietal attention network. Proc Natl Acad Sci USA, 110, 15806–15811. 10.1073/pnas.1313903110

74. Tang, Y.-Y., Hölzel, B.K., & Posner, M.I. (2015). The neuroscience of mindfulness meditation. Nature Reviews Neuroscience, 16, 213–225. 10.1038/nrn3916

75. Uddin, L.Q., Molnar-Szakacs, I., Zaidel, E., & Iacoboni, M. (2006). rTMS to the right inferior parietal lobule disrupts self-other discrimination. Soc Cogn Affect Neurosci, 1, 65–71. 10.1093/scan/nsl003

76. Uddin, L.Q., Yeo, B.T.T., & Spreng, R.N. (2019). Towards a universal taxonomy of macro-scale functional human brain networks. Brain Topogr, 32, 926–942. 10.1007/s10548-019-00744-6

77. Valk, S.L., Bernhardt, B.C., Trautwein, F.M., Böckler, A., Kanske, P., Guizard, N., Collins, D.L., & Singer, T. (2017). Structural plasticity of the social brain: Differential change after socio-affective and cognitive mental training. Science Advances, 3, e1700489. 10.1126/sciadv.1700489

78. Vallar, G., & Calzolari, E. (2018). Unilateral spatial neglect after posterior parietal damage. Handb Clin Neurol, 151, 287–312. 10.1016/B978-0-444-63622-5.00014-0

79. Vossel, S., Geng, J.J., & Fink, G.R. (2014). Dorsal and ventral attention systems: distinct neural circuits but collaborative roles. Neuroscientist, 20, 150–159. 10.1177/1073858413494269

80. Yordanova, J., Kolev, V., Mauro, F., Nicolardi, V., Simione, L., Calabrese, L., Malinowski, P., & Raffone, A. (2020). Common and distinct lateralised patterns of neural coupling during focused attention, open monitoring and loving kindness meditation. Sci Rep, 10, 7430. 10.1038/s41598-020-64324-6

81. Yordanova, J., Kolev, V., Nicolardi, V., Simione, L., Mauro, F., Garberi, P., Raffone, A., & Malinowski, P. (2021). Attentional and cognitive monitoring brain networks in long-term meditators depend on meditation states and expertise. Sci Rep, 11, 4909. 10.1038/s41598-021-84325-3

82. Yordanova, J., Kolev, V., Verleger, R., Heide, W., Grumbt, M., & Schürmann, M. (2017). Synchronization of fronto-parietal beta and theta networks as a signature of visual awareness in neglect. NeuroImage, 146, 341–354. 10.1016/j.neuroimage.2016.11.013

